# Identification of new growth regulators using cross-species network analysis in plants

**DOI:** 10.1101/2021.10.25.465753

**Authors:** Pasquale Luca Curci, Jie Zhang, Niklas Mähler, Carolin Seyfferth, Chanaka Mannapperuma, Tim Diels, Tom Van Hautegem, David Jonsen, Nathaniel Street, Torgeir R. Hvidsten, Magnus Hertzberg, Ove Nilsson, Dirk Inze, Hilde Nelissen, Klaas Vandepoele

## Abstract

With the need to increase plant productivity, one of the challenges plant scientists are facing is to identify genes playing a role in beneficial plant traits. Moreover, even when such genes are found, it is generally not trivial to transfer this knowledge about gene function across species to identify functional orthologs. Here, we focused on the leaf to study plant growth. First, we built leaf growth transcriptional networks in *Arabidopsis thaliana*, maize (*Zea mays*), and aspen (*Populus tremula*). Next, known growth regulators, here defined as genes that when mutated or ectopically expressed alter plant growth, together with cross-species conserved networks, were used as guides to predict novel Arabidopsis growth regulators. Using an in-depth literature screening, 34 out of 100 top predicted growth regulators were confirmed to affect leaf phenotype when mutated or overexpressed and thus represent novel potential growth regulators. Globally, these new growth regulators were involved in cell cycle, plant defense responses, gibberellin, auxin, and brassinosteroid signaling. Phenotypic characterization of loss-of-function lines confirmed two newly predicted growth regulators to be involved in leaf growth (*NPF6.4* and *LATE MERISTEM IDENTITY2*). In conclusion, the presented network approach offers an integrative cross-species strategy to identify new genes involved in plant growth and development.

**One-sentence summary:** Cross-species network analysis results in the identification and validation of new growth regulators in Arabidopsis.

## Introduction

The need to increase plant productivity reveals that, despite the detailed information gained on plant genomes, modelling plant growth and translating the molecular knowledge obtained in model plant species to crops is not trivial (Nuccio et al., 2018; Simmons et al., 2021, Inze and Nelissen, under review). Plant organ growth is one of the processes that is well-studied in model plants (Vercruysse et al., 2020a), playing a major role in affecting plant productivity (Sun et al., 2017). New plant organs are formed and then grow continuously throughout development. Upon adverse conditions, growth adjustments are among the first plant responses, rendering growth regulation an important yield component (Gray and Brady, 2016; Nowicka, 2019). The growth of plants involves complex mechanisms controlling processes from the cellular to the whole-organism level (Verbraeken et al., 2021). However, which growth zones or cell types are most important in controlling organ growth is not always clear.

Numerous genes, which we refer to as growth regulators, have been identified that when mutated or ectopically expressed alter organ size, such as leaf size, in plants. Detailed transcriptome and functional analyses have revealed that many of these genes are part of functional modules conserved across plant species (Vercruysse et al., 2020b). Previous research has shown that largely similar cellular and molecular pathways govern the fundamental growth processes in dicots and monocots (Anastasiou et al., 2007; Nelissen et al., 2016). This observation is based on the presence of functionally conserved orthologous growth regulators which promote organ growth in both dicots and monocots. Notable examples are genes encoding CYP78A, ARGOS, rate limiting GA biosynthesis enzymes, BRI1, ANGUSTIFOLIA3 and GROWTH-REGULATING FACTORS (Powell and Lenhard, 2012; Vercruysse et al., 2020a).

The complex and highly dynamic nature of the regulatory networks controlling complex traits makes the identification of new growth regulatory genes challenging (Baxter, 2020). Moreover, duplication events across the plant kingdom have caused a general enlargement of gene families and, with it, plant-and tissue-specific functional specialization (Jones and Vandepoele, 2020). It became clear that, even when the gene space is well characterized and conserved, the translation from model species to crops is not straightforward (Pan et al., accepted; Inze and Nelissen, in press). One of the bottlenecks lies in the complexity of crop genomes, such as polyploidy, and the subsequent difficulty to identify functional orthologs.

Gene orthology information is essential to transfer functional annotations from model plants with high-quality annotations (e.g. *Arabidopsis thaliana*) to other species. Functional annotations derived from experimental evidence can be used to identify relevant orthologs and drive gene function discovery in crops (Lee et al., 2015, 2019). This approach is not straightforward, mainly for two reasons: first, the orthology approach normally leads to the identification of complex (one-to-one, one-to-many and many-to-many) orthology relationships (Movahedi et al., 2011; Van Bel et al., 2012); second, for genes with multiple orthologs, it has been observed that the closest ortholog in terms of protein sequence similarity is often not the closest ortholog in terms of regulation, indicating that identifying functionally conserved orthologs is challenging (Patel et al., 2012; Netotea et al., 2014).

Biological networks offer the means to study the complex organization of gene interactions. Densely connected network clusters form gene modules, defined as groups of linked genes with similar expression profiles (i.e. co-expressed genes), which also tend to be co-regulated and functionally related (Heyndrickx and Vandepoele, 2012; Klie et al., 2012). Although transferring network links from better annotated species to crops is the most intuitive approach and has proven to be helpful (Ficklin and Feltus, 2011; Obertello et al., 2015), it has been shown that only ∼20-40% of the co-expression links are conserved in pairwise comparison of *Arabidopsis thaliana* (Arabidopsis), *Populus*, and *Oryza sativa* (Netotea et al., 2014). On the other hand, it has been shown that using gene modules that are conserved across species can increase the amount of biological knowledge transferred from one species to another (Mutwil et al., 2011; Heyndrickx and Vandepoele, 2012; Cheng et al., 2021). Such conserved gene modules mirror biological processes conserved across species, meaning that the orthologous genes present in these modules are involved in the same process and potentially perform the same function (Ruprecht et al., 2011; Stuart et al., 2003). Significantly conserved cross-species modules (with many shared orthologs) can be used to transfer gene function annotations and analyze expression conservation for paralogs involved in complex many-to-many orthology relationships. A guilt-by-association approach can also then be used to infer functions of unknown genes from the functions of co-expressed annotated genes (Wolfe et al., 2005; Lee et al., 2010; De Smet and Marchal, 2010; Klie et al., 2012; Rhee and Mutwil, 2014).

Here, we aimed at developing an integrative approach to identify functionally conserved regulators, leveraging high-resolution transcriptomes and the power of cross-species network biology. In particular, we chose leaf as a system to study plant growth, as high-quality datasets covering cell proliferation and expansion are available in three plant species: two dicotyledonous plants, the annual plant Arabidopsis and the perennial plant *Populus tremula* (aspen), and one monocotyledonous plant, *Zea mays* (maize). We leveraged these data to construct aggregated gene networks for each species and identified, through gene neighborhood conservation analysis, genes with cross-species network conservation. Subsequently, we used known plant growth regulators, belonging to various functional modules and influencing growth of different plant organs, as guide genes to predict new putative growth regulators among these conserved genes. For the top 100 predicted growth regulators, we screened the literature to investigate if predictions linked to leaf growth were obtained. For a subset of highly ranked predictions with no reported information on plant growth, we performed phenotypic analyses and succeeded to validate two novel Arabidopsis growth regulators.

## Results

### Network construction and gene neighborhood conservation analysis

To perform network construction based on gene expression information, we used transcriptomic data from leaves, which were selected as a representative system to study plant growth. This choice was primarily motivated by the well-known similarities in leaf growth regulation across dicots and monocots, which make cross-species comparison of gene networks straightforward and useful for gene function discovery (Vercruysse et al., 2020b). Secondly, our motivation relied on the availability of large-scale expression profiling studies, which allow selecting similar samples and constructing a congruent dataset for the different species. Expression compendia were built for Arabidopsis, maize and aspen that contained a minimum of 24 leaf samples (Figure 1, step 1; Supplemental Table 1; Supplemental Methods). These expression compendia all include developmental stages with active cell proliferation and cell expansion. The Arabidopsis expression compendium was composed of three main developmental phases: cell proliferation, cell expansion and the transition between these two phases. For maize, the developmental expression compendium included a newly generated high-resolution dataset and covered cell proliferation, cell expansion and mature phases of development (Supplemental Methods). For aspen, samples covered the developmental stages ranging from the very youngest leaf primordia to fully expanded and mature leaves. In total, expression data covered 20,313 genes for Arabidopsis, 29,383 genes for maize, and 35,309 genes for aspen (Dataset 1).

**Figure 1.**
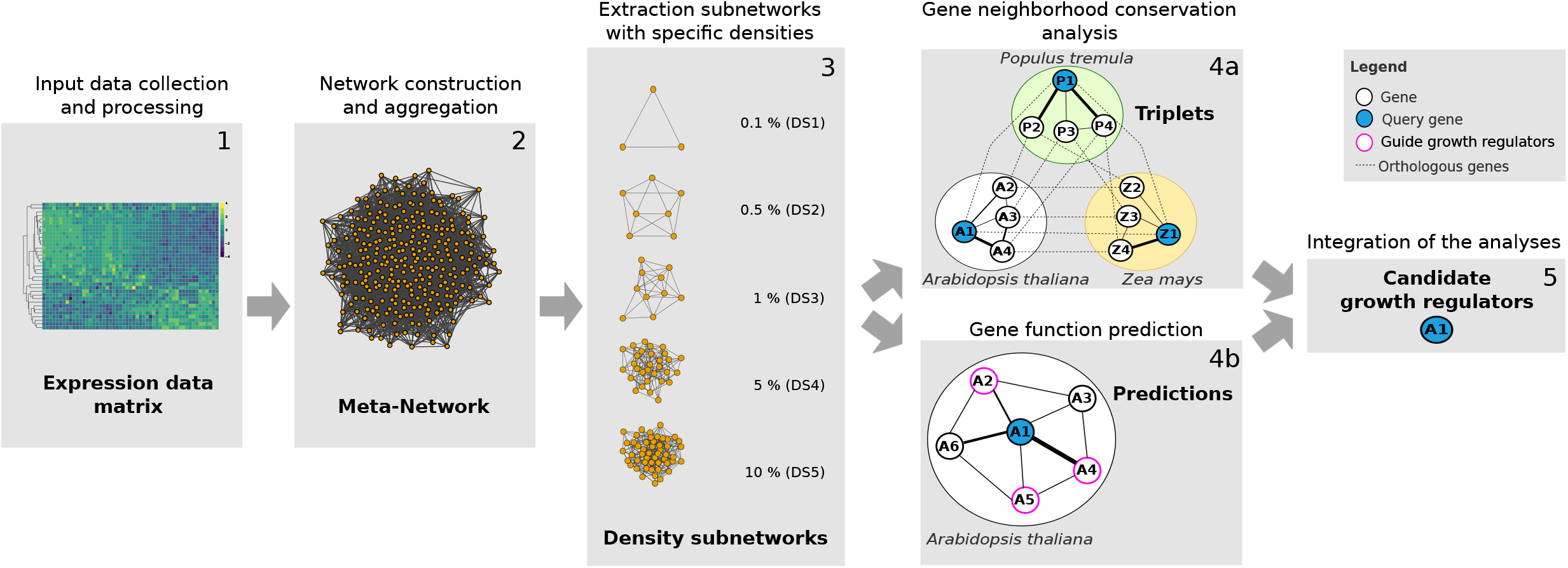
Outline of the cross-species network approach to identify new candidate growth regulators. For Arabidopsis, maize and aspen, the expression data (step 1) is used as input to construct a fully connected meta-network per species (step 2). Subsequently each meta-network is split into five density subnetworks (DSs) by applying specific density cutoffs (step 3). These DSs are the input for two different analyses: they are used first as input to compute cross-species gene neighborhood conservation (step 4a). Secondly, they are used to predict new functions via guilt-by-association (step 4b). This leads to gene function annotations of query genes (blue circles) based on prior knowledge on growth regulators (purple circles). Edge thickness defines in which subnetwork the interaction is conserved (line thickness represents the DS and ranges from 1, the most stringent DS represented by the thickest line, to 5, the least stringent DS represented by the thinnest line). Finally, the results of these two analyses (steps 4a and 4b) are integrated to obtain a list of GR candidates (step 5).

The network construction was performed for each species with Seidr, a toolkit to perform multiple gene network inferences and combine their results into a unified meta-network (Schiffthaler et al., 2018). For each network inference algorithm included, a fully connected weighted gene network was constructed. These were in turn aggregated into a weighted meta-network (simply “network” hereinafter, Figure 1, step 2). When applying a weight threshold, the network density was defined as the ratio between the number of links with a weight higher than this threshold and the number of links in the weighted network. To dissect the network structure, several thresholds were used to subset the networks into more stringent density subnetworks (DSs). For each species network, five DSs were obtained ranging from DS1 (top 0.1% links) with an average of 358,455 links, to DS5 (top 10% links) with an average of 35,845,512 links (Figure 1, step 3), with higher densities corresponding to a higher number of neighbors for each gene in the network (Supplemental Figure 1). A gene’s neighborhood is defined as all genes connected with this gene for a given network.

Genes showing gene neighborhood conservation across species are part of conserved functional modules controlling distinct biological processes. This implies that the conserved network containing these genes confers a selective advantage and therefore that these genes are functionally related (Stuart et al., 2003). However, which gene neighborhood size to select to identify conserved growth-related functional modules is not straightforward, as being too stringent might lead to the loss of valuable interactions while being too relaxed might include non-functional interactions potentially representing noise (Movahedi et al. 2012). To identify genes showing network conservation in different species, a gene neighborhood conservation analysis was performed using each DS and the information on the orthology relationships between Arabidopsis, maize and aspen genes (Figure 1, step 4a). The network neighborhood of a gene is represented by all genes connected to it, at a given threshold. This concept was used to identify “triplets” (Dataset 2), each containing three orthologous genes across Arabidopsis, maize and aspen with statistically significant overlaps between their gene network neighborhoods (see Methods). In an example triplet (Figure 1, step 4a), a specific Arabidopsis gene *A1*, will have an ortholog *Z1* in maize and another ortholog *P1* in aspen and these three genes will have a significant overlap of their gene network neighborhoods. Due to the complex orthology relationships that exist in plants, each gene can belong to one or multiple triplets as it can have one or more orthologs. For example, an Arabidopsis gene with only one ortholog in maize and aspen, assuming they have significant overlap of their gene network neighborhoods, will belong to one triplet. In contrast, another Arabidopsis gene with two orthologs in maize and three in aspen, assuming they also all have significant overlaps of their gene network neighborhoods, will belong to six triplets. We refer to the set of unique genes that are part of triplets as “triplet genes”. Next, the conserved gene neighborhoods were used to dissect the complex network structures of these plants and to functionally harness the orthology relationships. The cross-species networks are available in an interactive web application (https://beta-complex.plantgenie.org).

### Delineation of conserved growth regulators

Since the output of cell proliferation and expansion are strongly contributing to leaf size, we hypothesized that the generated triplets were an excellent source to extract orthologs potentially altering plant growth, representing conserved GRs. Growth regulators typically act by stimulating cell proliferation (yielding a higher cell number, as in the case of GRF (GROWTH-REGULATING FACTOR) and GIF (GRF-INTERACTING FACTOR) proteins (Lee et al., 2009)) and/or cell expansion (as in the case of *ZHD5* (*ZINC-FINGER HOMEODOMAIN 5*) (Hong et al., 2011)). We generated a list of known GRs (“primary-GRs”) covering 71 primary-GRs from Arabidopsis, 71 from aspen and eight from maize. While the Arabidopsis and maize GRs mainly have a role in controlling leaf size, the aspen GRs are affecting stem size. In both organs, cell proliferation and expansion play an important role in controlling growth (Serrano-Mislata and Sablowski, 2018). This list of genes was obtained by collecting scientific literature and by phenotypic analysis of mutant and over-expression lines in Arabidopsis, maize, and aspen. We then used the triplets to transfer GRs from maize and aspen to Arabidopsis (“translated-GRs”). In other words, primary-GRs from maize and aspen, also identified as triplet genes, were used to extract Arabidopsis orthologs with gene neighborhood conservation. The primary-GRs and translated-GRs were finally merged and filtered for high expression variation in the Arabidopsis expression compendium to retain only those active during either cell proliferation or cell expansion. The resulting set, named “expression-supported GRs” (Supplemental Table 2, Supplemental Figure 2), was composed of 82 GRs, including 24 Arabidopsis primary-GRs and 58 translated-GRs (*GRF2* and *GA20OX1* (*GIBBERELLIN 20-OXIDASE 1*) were shared between primary-GR and translated-GR sets). According to their expression profiles in Arabidopsis, 35 expression-supported GRs showed maximal expression during cell proliferation, including several proliferation marker genes like GROWTH-REGULATING FACTORs (e.g. *GRF1*, *GRF2*, *GRF3*), AINTEGUMENTA (*ANT* (Mizukami and Fischer, 2000) and *KLUH* (Anastasiou et al., 2007)), and 47 expression-supported GRs had increased expression during cell expansion, such as *GA20Ox1* (Barboza et al., 2013) and *BR ENHANCED EXPRESSION 2* (*BEE2* (Friedrichsen et al., 2002)).

The 82 expression-supported GRs (from here on simply referred to as “GRs”) represent our guide genes, obtained by the integration of prior knowledge on plant growth and the cross-species gene neighborhood conservation approach, to identify new candidate GRs.

### Functional analysis of cross-species conserved networks underlying leaf cell proliferation and expansion

To explore cross-species conserved genes that function during cell proliferation and expansion, we performed a Gene Ontology (GO (Ashburner et al., 2000)) functional enrichment analysis of the Arabidopsis triplet genes from each DS across two sets: (1) all triplet genes (All) and (2) the subset of triplet genes including the 82 GRs and their co-expressed triplet genes (Growth regulator-related triplet genes) (Figure 2). The total number of triplets ranged from 1,739 (DS1) to 243,645 (DS5) (Figure 2A; Dataset 2). To assess the significance of these numbers, a permutation approach was employed where the orthology relationships were randomized 500 times and the number of triplets obtained from each permutation was recorded. The number of triplets observed were highly significant with not a single permutation for any DS exceeding the number of triplets observed in the non-permuted data (p-value<0.002). The number of unique Arabidopsis triplet genes ranged from 211 (DS1) to 6,526 (DS5) indicating that less sparse networks tend to have more genes and more conserved gene neighborhoods (Figure 2A). Interestingly, GRs and their network neighbors on average made up 71% of the triplet genes across the five DSs, suggesting that leaf growth-related gene networks are well conserved during leaf development across plant species. For simplicity, from here on we will refer to triplet genes at a specific DS as, for example at DS1, “genes conserved at DS1”. The functional enrichment (Figure 2B) showed that triplet genes from the most stringent subnetwork (DS1) were enriched for basal biological processes during leaf development, including photosynthesis (e.g. glucose metabolic process, response to light and carbon fixation) and translation (e.g. large and small ribosomal subunits). Processes such as cell division and cell cycle regulation were significantly enriched for genes conserved at DS2 and DS3, including genes coding for cyclins (type A, B, D and P), cyclin dependent kinases (*CDK*) and their subunits (*CKS*), and other genes involved in the spindle formation (i.e. *MICROTUBULE-ASSOCIATED PROTEINS (MAP)65-4 and −5*). Cell expansion-related processes were identified among genes conserved at DS3 and included genes coding for expansins (EXP) and xyloglucan endotransglucosylases/hydrolases (XTH). Genes conserved at the two least stringent subnetworks (DS4 and DS5) were enriched for GO terms related to cell wall organization (e.g. lignan biosynthesis, pectin degradation, lignin metabolism), defense response to biotic and abiotic stresses (e.g. defense response to oomycetes, response to salt stress and heat stress), and transmembrane transport and hormone signaling (e.g. response to auxin, ethylene and brassinosteroid). The category “regulation of transcription” was enriched for genes conserved at DS3, DS4, and DS5. GRs were significantly over-represented in subnetworks starting from DS2, indicating that GRs have highly conserved gene network neighborhoods. Most of the GRs (87%) were conserved in one or more DSs (Figure 2C).

**Figure 2.**
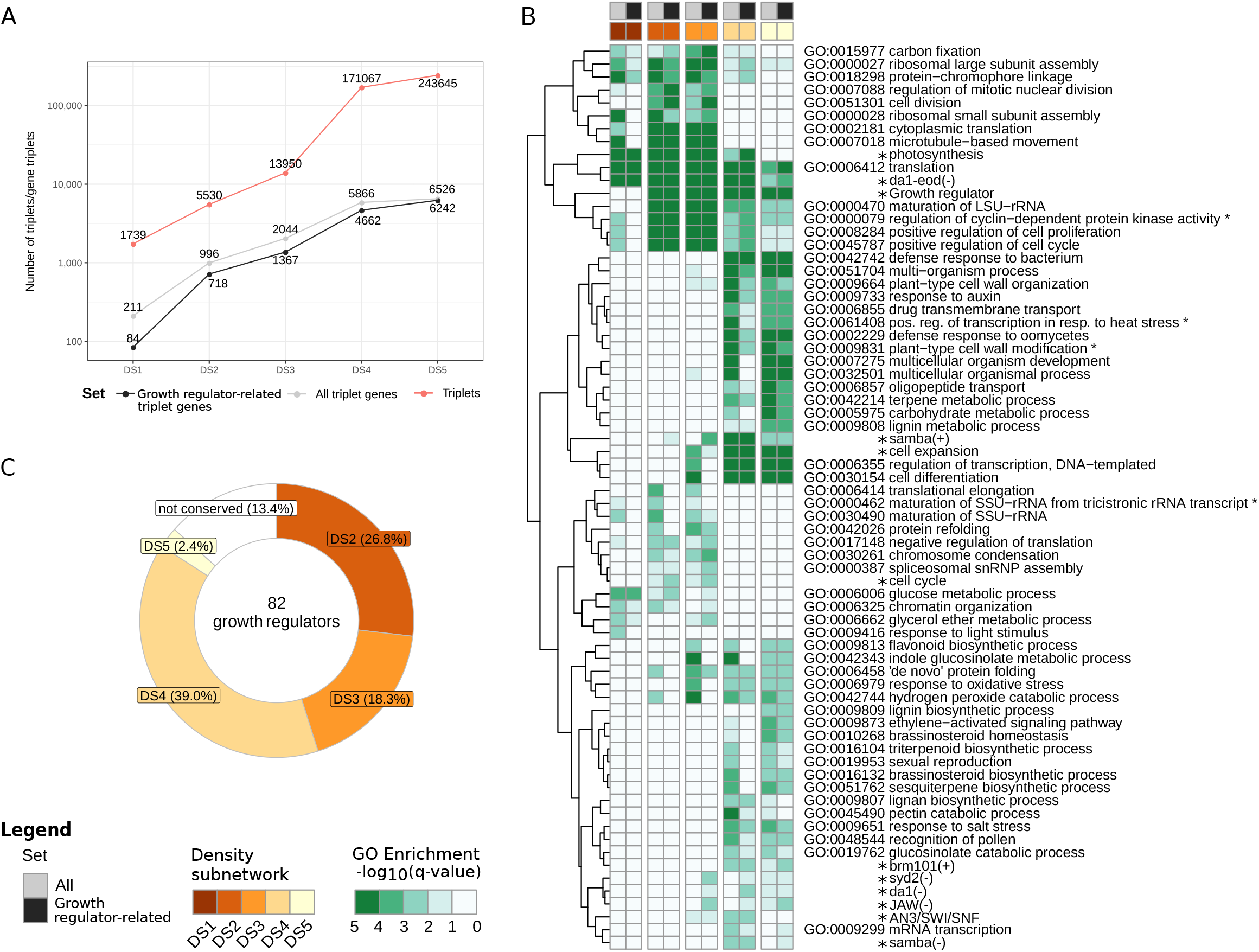
Triplets and their functional enrichments in cross-species conserved leaf networks. (A) The number of triplet genes showing cross-species neighborhood conservation is plotted for all density subnetworks (DSs). (B) The functional over-representation of biological processes of interest is summarized for two sets: all triplet genes (All) and for growth regulators and their network neighbor (Growth regulator-related) triplet genes, subset of all triplet genes, in each DS. Functional categories marked with asterisks (*) belong to leaf growth modules described in Vercruysse et al. (2020) and to the differentially expressed gene sets from relevant studies on plant development (Bezhani et al., 2007; Gonzalez et al., 2010; Eloy et al., 2012; Vercruyssen et al., 2014; Vanhaeren et al., 2017). (C) Overview of growth regulators with (and without) cross-species neighborhood conservation in different DSs.

Among the GRs conserved at DS2, 32% were transcription factors (TFs), including regulators of cell cycle (e.g. *AINTEGUMENTA*) and cell elongation such as *BEE2* and its homolog *HBI1* (Supplemental Figure 3). These results suggest a conserved role of these TFs in leaf development across the three plant species. Genes involved in hormone-mediated transcriptional regulation (*INDOLEACETIC ACID-INDUCED PROTEIN (IAA)3*, *IAA14*, *IAA30*, and *AUXIN RESISTANT* (*AUX*)*1*) were also detected. Cell growth regulators, including the GRF family, were found conserved and, among them, *GRF2* was conserved at DS2. Literature information on differentially expressed gene (DEG) sets from perturbation experiments was also included in the functional enrichment analyses for several primary-GRs. In particular, genes up- and down-regulated in *SAMBA* loss-of-function mutants (Eloy et al., 2012) and *JAW (JAGGED AND WAVY)* overexpression lines (Gonzalez et al., 2010) were significantly enriched in the GR-related set (Figure 2B). Whereas *SAMBA* plays a key role in organ size control (seeds, leaves and roots), transgenic overexpression lines of *JAW* showed enlarged leaves and an increased cell number, indicative of prolonged cell proliferation (Gonzalez et al., 2010; Eloy et al., 2012). An additional functional enrichment analysis was performed focusing on TF families to identify their cross-species conservation level. In particular, genes conserved from DS2 to DS5 (Supplemental Figure 4) were significantly enriched for the ETHYLENE RESPONSE FACTOR (ERF) family (q-value < 0.01), which has a recognized role in plant growth (Dubois et al., 2018). At DS3, among others, MYB and WRKY TF families, known to be involved in developmental processes, appeared strongly conserved. At the least stringent DSs (DS4 and DS5) we could observe other conserved TF families like DOF (regulating the transcriptional machinery in plant cells), MIKC-MADS (involved in floral development) and NAC (with functions in plant growth, development and stress responses) (Lehti-Shiu et al., 2017). For TFs conserved at DS2, a significant enrichment was observed for the CONSTANS-like TF-family when considering GR-related triplet genes and included *BBX3*, *BBX4, BBX14* and *BBX16*. A number of BBX proteins have been linked with photomorphogenesis, neighborhood detection, and photoperiodic regulation of flowering (Vaishak et al., 2019).

### Network-based prediction of novel growth regulators

Apart from analyzing the conservation level of known GRs, we subsequently investigated if new GRs could be identified. To obtain high-quality GR predictions, a combined strategy was adopted to leverage the known GRs and the gene neighborhood conservation analysis through a guilt-by-association (GBA) approach. The GBA principle states that genes with related function tend to be protein interaction partners or share features such as expression patterns or close network neighborhood (Oliver Stephen, 2000). First, gene function prediction through GBA was performed, where the known GRs were used as guide genes for network-based gene function discovery (Figure 1, step 4b). New gene functions were assigned through functional enrichment in the Arabidopsis networks, at different DSs. As a result, genes that were part of network neighborhoods significantly enriched for guide GRs were classified as newly predicted GRs, and a GBA score was assigned to quantify the strength of the predicted GRs (see Methods). Secondly, the new predictions (Figure 1, step 4b) were filtered for those already identified as triplet genes (Figure 1, step 4a). These filtered predictions (Figure 1, step 5), forming the predicted GR set, were labelled with their species names if they were part of the guide GRs (primary or translated-GR) or with “new” if they were novel (Supplemental Table 3). This approach led to 2206 GR predictions, of which 66 were guide GRs. For the latter, 11 were uniquely from the Arabidopsis GR primary set, 53 uniquely from the aspen translated-GRs, and the remaining two were shared among species. Note that the recovery of known GR genes would be zero in case the network would be random and not capture growth-related transcriptional information. From DS1 to DS5, the subsets of GR predictions covered 175, 496, 421, 891 and 223 genes, respectively (Supplemental Table 3). Overall, the biological processes observed for the conserved predictions agreed with those observed for all triplet genes (Figure 2).

To evaluate the reliability of the predicted GR set and its potential use for discovering genes with a significant effect on plant growth, the public phenotype database RARGE II (Akiyama et al., 2014), covering 17,808 genes and 35,594 lines, was screened obtaining a list of 391 Arabidopsis genes that, if mutated, caused a phenotype change in Arabidopsis leaf length, width and/or size (RARGE II leaf trait genes, Supplemental Table 4). When investigating the gene recovery for the RARGE II leaf trait genes (Figure 3), a clear trend was observed in phenotype recovery ranging from DS1, with higher recovery (∼3 and ∼4.3 fold enrichment compared to what is expected by chance for proliferation and expansion, respectively), to DS5, with almost no recovery. This result indicates that, among all DSs, DS5 is the least suitable one to identify genes with a potential effect on leaf phenotype.

**Figure 3.**
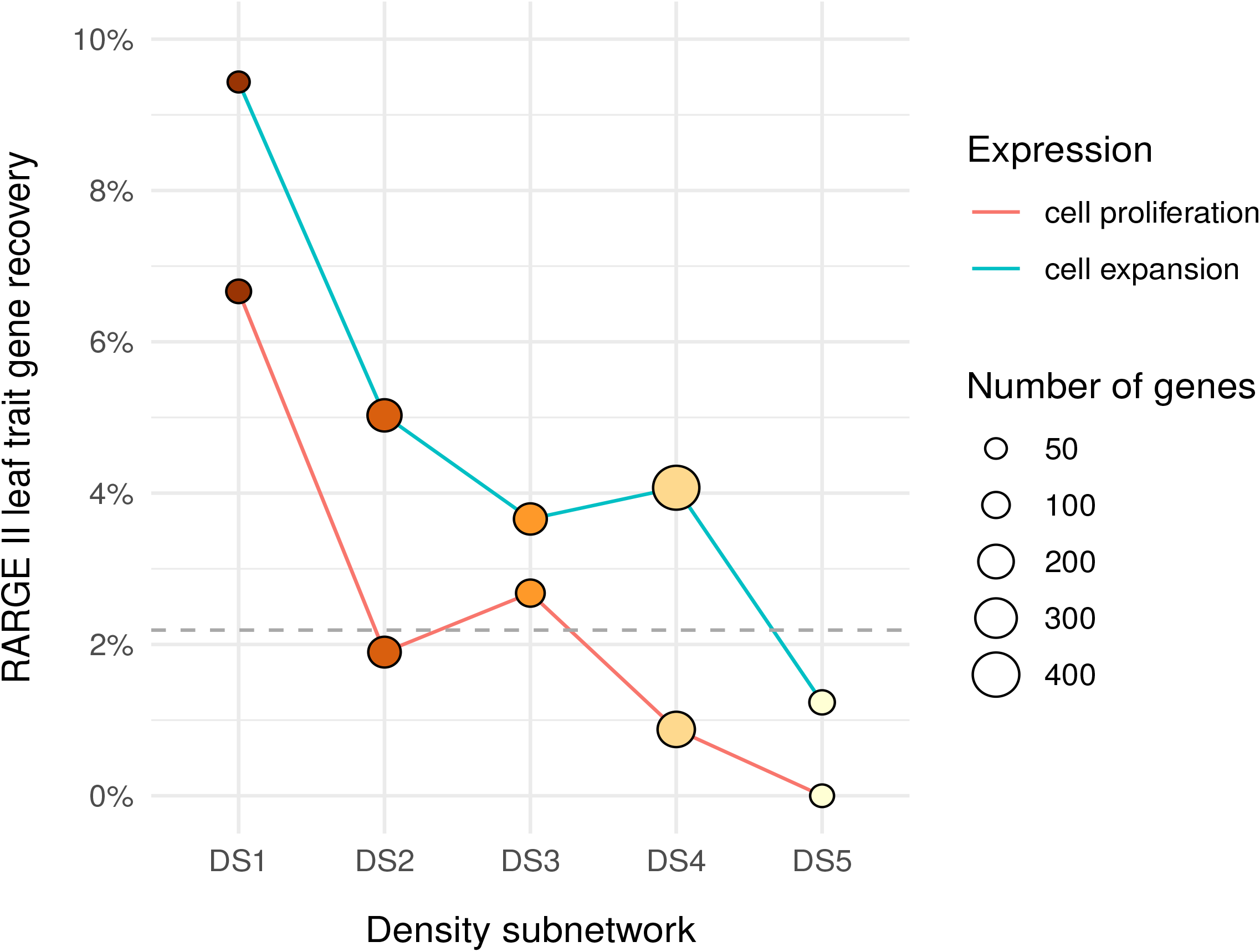
Recovery of RARGE II leaf trait genes for each DS split in proliferation and expansion. The grey dashed line indicates the leaf-related phenotype gene recovery expected by chance (within the RARGE II dataset).

### Validation of GR predictions using literature and leaf phenotyping

To validate the assumption that the GR predictions top ranked by GBA are more likely to show a plant growth-related phenotype, an in-depth literature analysis was performed to summarize the connection with different growth-related pathways (Supplemental Table 5) and to score known growth-related phenotypes for the top 100 GR predictions (Supplemental Table 6). For 61 of these 100 predicted genes, mutant lines and/or lines with ectopic expression were reported. For 34 out of the 61 genes (55.7%), obvious alterations to leaf size and shape as well as petiole length were reported when mutated or overexpressed (Supplemental Table 6).

Functional analysis of the 34 genes with described leaf phenotypes revealed their involvement in several biological processes and pathways such as cell cycle regulation, hormone response, photosynthesis, carbon utilization and cell wall modification (Figure 4). Importantly, we could find conserved relationships between five specific genes active in the expansion phase: CATIONIC AMINO ACID TRANSPORTER *(CAT)2*, *THIOREDOXIN* X *(THX)*, BETA CARBONIC ANHYDRASE *(BCA)4*, *CA2*, and *PMDH2*. Among them, *CAT2* and *BCA4* were also high ranked by GBA score. For the proliferation cluster, we could observe strong relationships between *ANT*, *OBP1*, *GRF2*, *CYCD3;3*, *GL1*, *HTA8 (HISTONE H2A 8)*, and *AN3*. Among them, we identified TFs mainly involved in cell cycle process (*ANT*, *OBP1*, *GRF2*), cell wall (*GL1*), and hormone signaling pathways such as jasmonate (*GL1*), abscisic acid (*ANT*), and gibberellin (*GL1*). Twenty-seven of the 61 predictions with knock-down mutations and/or ectopic expression lines did not show a correlation with leaf growth, which may be partially due to the redundancy of large gene families or that the leaf phenotype was not explored in those studies. Additionally, three of these 27 genes have been reported to influence root or hypocotyl development, which may also contribute to overall plant growth and organ size.

**Figure 4.**
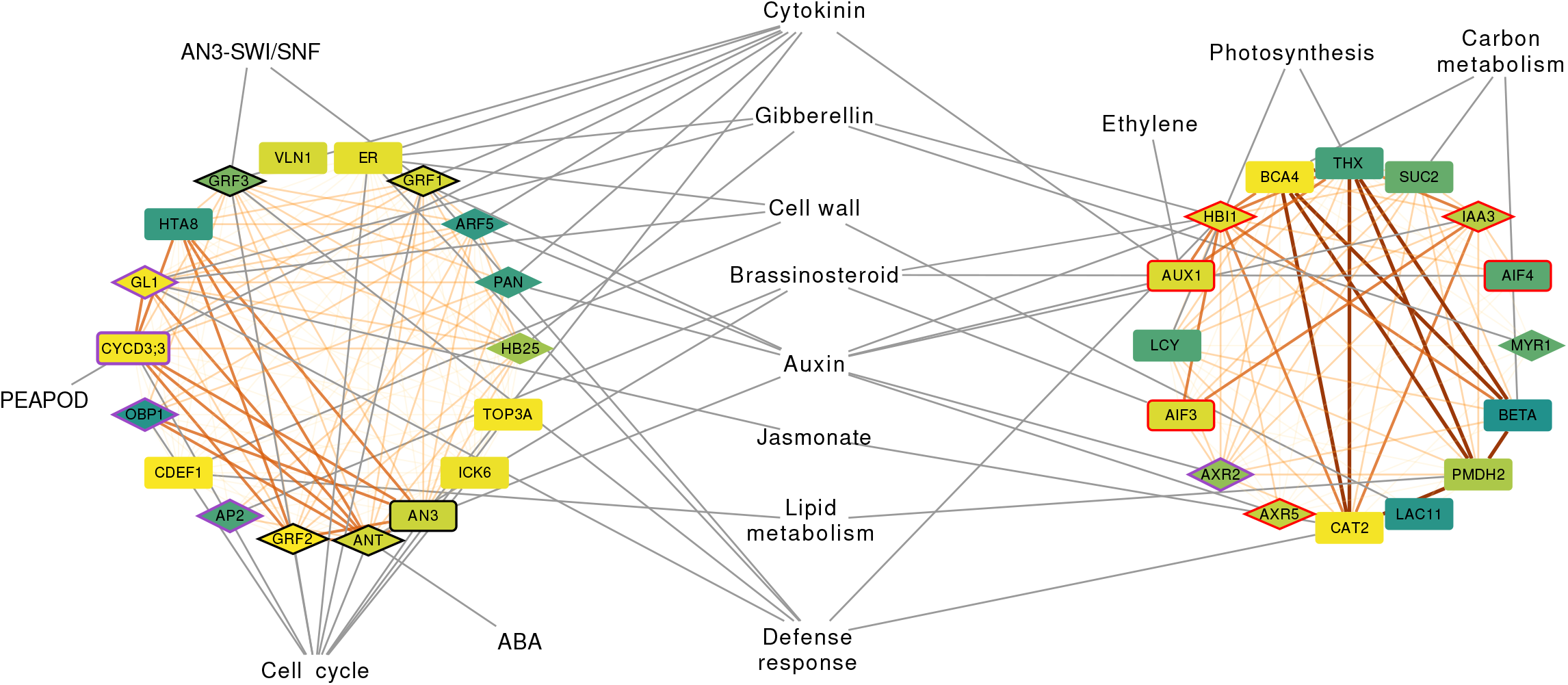
Gene-function network of the 34 phenotype-related genes out of the top 100 predicted growth regulators. Predictions are clustered by expression profile (proliferation on the left and expansion on the right). Node label colours from yellow (strong) to dark green (weak) represent the reliability of the gene prediction (GBA score). Node border colours indicate known growth regulators from Arabidopsis (black), known growth regulators from aspen (red), and Arabidopsis known growth regulators paralogs (violet). Diamonds represent transcription factors. Links from dark orange thick (DS1) to light orange thin (DS5) represent the density subnetwork where the genes were found connected. Genes are linked with their respective growth-related pathways (centered if connecting to both proliferation and expansion related genes) by grey links. Anti-correlation links (connecting proliferation with expansion genes) were removed for clarity.

To further validate the role of these new candidate GRs in the leaf development, the system that we chose to study plant growth, we collected the mutants of nine genes among the 27 predicted GRs which have not been reported with a leaf phenotype (Supplemental Table 7). Molecular identification of these mutants was conducted and a detailed analysis of leaf growth in controlled long-day soil-grown conditions was made (Supplemental Figure 5). By following the projected rosette area (PRA), compactness and stockiness of each mutant line over time, this phenotypic characterization revealed that the mutants of two GR candidate genes showed altered rosette growth. The mutant lines of a putative nitrate transporter gene *NPF6.4/NRT1.3*, *sper3-1* and *sper3-3*, both displayed decreased PRA compared with the wild-type plants (Figure 5A). The *sper3-1* harbored a mutation at a conserved glutamate *of NRT1.3,* while the T-DNA line *sper3-3* was a knockout allele (Tong et al., 2016). The reduction in size of *sper3-3* was smaller and occurred later in development compared with *sper3-1*. Before bolting (26 DAS), *sper3-1* and *sper3-3* were 37.3% and 13.2% smaller, respectively, compared with the wild-type (Supplemental Table 7). Both *sper3-1* and *sper3-3* showed significantly reduced leaf number compared to wild type (Figure 5, Supplemental Figure 6). Besides *NPF6.4*, the mutants of *LATE MERISTEM IDENTITY2* (*LMI2*) which has been reported to be required for correct timing of the meristem identity transition (Pastore et al., 2011), also showed altered rosette growth. In standard long-day conditions in soil, a significant reduction of PRA was detected in *lmi2-1*, which displayed elevated *LMI2* expression in seedlings. By contrast, the *lmi2-2* mutants in which the T-DNA insertion gave rise to a truncated non-functional LMI2 protein, exhibited significantly increased PRA and were 13.5% larger than the wild-type plants at 26 DAS (Figure 5B and Supplemental Table 7). Among *LMI2 mutants, lmi2-2* showed significantly increased leaf number (Figure 5, Supplemental Figure 6). Both *NPF6.4* and *LMI2* were highly ranked by GBA (rank 18 and 20, respectively), which further implies that the predictions with a low GBA score are more likely to show a leaf phenotype. Although the leaf was the model system chosen and analyzed in this study, we do not exclude that the predicted candidate GRs, including the validated *NPF6.4* and *LMI2*, might also alter the growth of other organs. Taken together, these experimentally validated genes lend additional support to the potential of our predictions for plant growth regulation.

**Figure 5.**
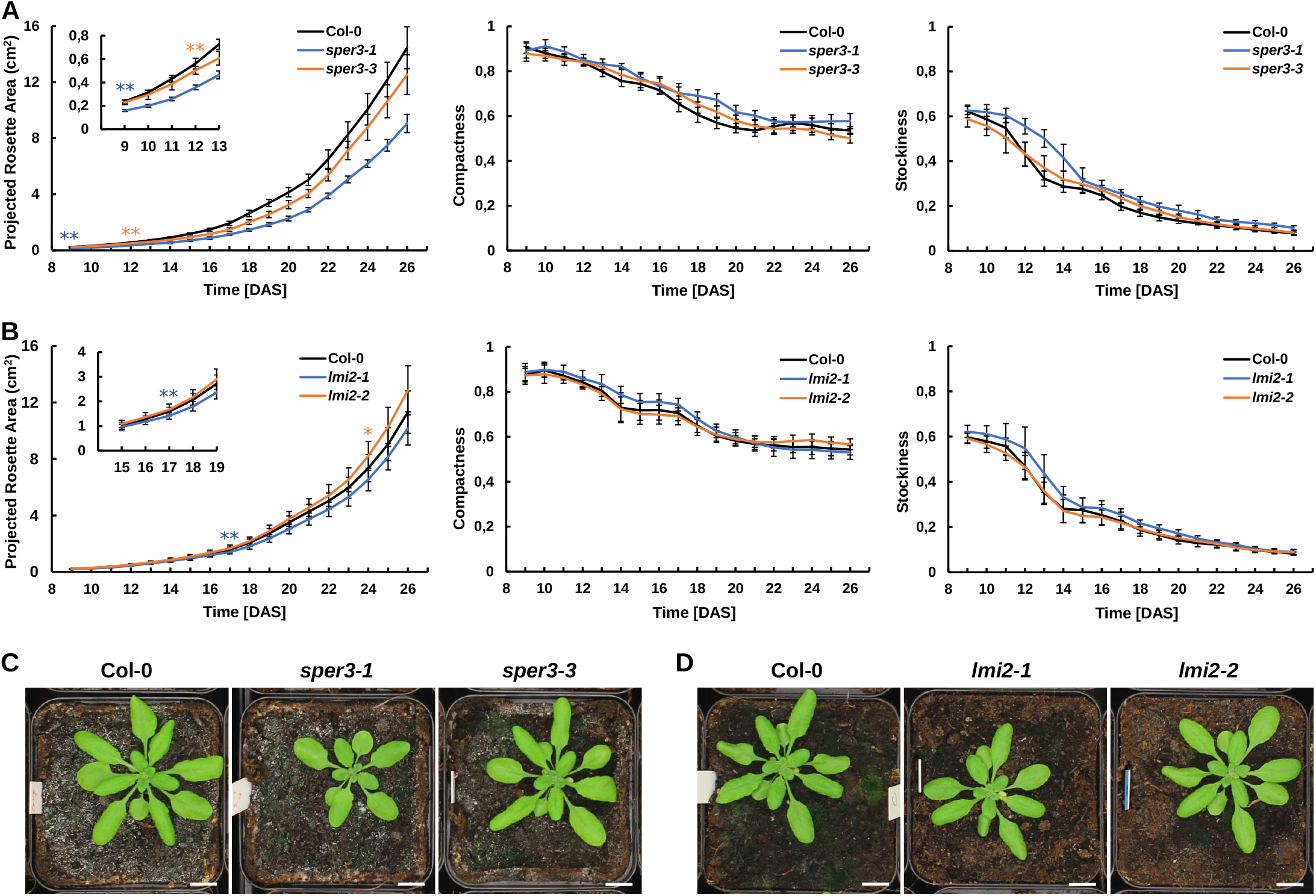
Mutants of predicted growth regulators *NRT1.3* and *LMI2* showed altered rosette growth. (A-B) Dynamic growth analysis of projected rosette area, compactness and stockiness over time of wild-type Col-0 and the mutants of *NRT1.3* (A) and *LMI2* (B) in soil. The asterisks represent the time points at which differences in the PRA become significant between the mutants and wild-type, as determined by Student’s t test (*, P<0.05; **, P<0.01). (C-D) Phenotype of 26-day-old mutants of *NRT1.3* (C) and *LMI2* (D). Scale bar = 1 cm.

## Discussion

In this study, we developed an integrative approach to identify candidate genes responsible for altering plant growth. To accomplish this, we used cross-species gene network analysis focusing on the leaf, given its similarities between dicots and monocots (Nelissen et al., 2016). To identify relevant context-specific gene interactions, it is highly recommended to focus the gene network analysis on a specific condition or context, rather than integrating multiple conditions (e.g. different stresses, growth conditions, development stages) (Pavlidis and Gillis, 2012; Liseron-Monfils and Ware, 2015; Serin et al., 2016). For this reason, expression datasets were generated and compiled capturing two main features of leaf growth: cell proliferation and cell expansion. These two processes are governed by similar cellular and molecular pathways across monocots and dicots (Nelissen et al., 2016), which inspired the selection of transcriptional datasets from two dicots (Arabidopsis and aspen) and one monocot (maize). The network construction was carried out integrating multiple inference methods to leverage the power and complementarity of different network inference algorithms (Marbach et al., 2012; Schiffthaler et al., 2018). To evaluate the strength of different biological signals in our network, the gene interactions, obtained after applying different network density cutoffs (DS1-5), were studied. Given that thousands of genes are expressed during leaf development, prioritizing new growth regulators starting from different developmental expression datasets is a major challenge. To do so, we relied on two main approaches: the guilt-by-association principle, which is frequently used for gene discovery, and network neighborhood conservation analysis, which detects significantly overlapping network neighborhoods across species to identify reliable functional orthologs (Movahedi et al., 2011; Netotea et al., 2014).

From the gene neighborhood conservation analysis on five different density subnetworks, we observed that, with an increased network density, the number of genes with conserved network neighborhood also grew. This is expected and is probably due to a greater statistical power when comparing larger neighborhoods (Netotea et al., 2014). Overall, as previously observed (Vercruysse et al., 2020b), the integration of different sequence-based orthology detection methods was important because of their complementarity, highlighting complex orthology relationships and evaluating the strength of the orthology support. Overall, 36% of the Arabidopsis genes (7,320 out of 20,313 genes present in the network) had conserved neighborhoods across Arabidopsis, aspen, and maize, in any of the five density subnetworks. This result is similar to what has been found across Arabidopsis, poplar and rice, although a different network construction pipeline was used there (Netotea et al., 2014).

From a plant breeding perspective, we were interested in cross-species functionally conserved predictions with experimental evidence in more than one species. *GA20-oxidase1* represents a well-known example of a GR that is functionally conserved across monocots and dicots. This gene was confirmed in our analyses to be conserved at the network neighborhood level. *GA20-oxidase1* is in fact a rate limiting enzyme for gibberellin growth hormone biosynthesis in Arabidopsis, aspen, maize and rice (Gonzalez et al., 2010; Nelissen et al., 2012; Qin et al., 2013; Eriksson et al., 2000). To validate the functional relevance of the predicted GRs, we screened the top 100 GR predictions and observed that, among the 34 Arabidopsis predicted genes with a known leaf phenotype in Arabidopsis, six were also already known to affect plant growth in aspen (here stem size). This result is not unexpected as overlapping regulatory mechanisms and genes are shared between primary and secondary meristems, which are responsible for the formation of plant tissues and organs (Baucher et al., 2007). The six translated-GRs were *AUX1*, *IAA3*/*SHY2*, *AXR5*, *AIF3*, *AIF4*, and *HBI1* and their expression in Arabidopsis was peaking at the cell expansion phase. The first three genes are auxin-related genes. Auxin is important for regulating root meristem growth and is crucial for root initiation and lateral root number. *AUX1* was translated from *aspen* Potra002054g16021 while *IAA3*/*SHY2* and *AXR5* were translated from aspen Potra000605g04596. For both these aspen genes, generated aspen RNAi lines exhibited an increase in stem size, an important indicator for tree biomass yield, connecting back to the underlying regulatory processes in the meristematic tissues (Supplemental Table 2). *AUX1* is an auxin transport protein which regulates auxin distribution across source (young leaf) and sink organs (young roots) (Marchant et al., 2002). *IAA3*/*SHY2* is crucial for root meristem development in Arabidopsis, being the converging point of cytokinin and auxin regulatory circuit (Li et al., 2020). Arabidopsis mutants for *AUX1* and *IAA3*/*SHY2* showed alterations in number and size of lateral roots (Tian and Reed, 1999; Marchant et al., 2002) while *AXR5* is an auxin response factor and mutant plants for this gene are resistant to auxin and show alterations of root and shoot tropisms (Yang et al., 2004). Our network results and phenotypes in aspen and Arabidopsis indicate that these genes also play an import role in meristem growth in other organs apart from root. *HBI1*, *AIF3*, and *AIF4*, encode a tier of interacting bHLH transcription factors downstream of BR and regulate the cell elongation in leaf blade and petiole (Bai et al., 2013; Ikeda et al., 2013). *AIF3* and *AIF4* were translated from Potra004144g24626 while *HBI1* was translated from Potra186144g28414. These two aspen genes have been tested with an overexpression approach in aspen trees showing even a bigger increase in stem size as compared with the auxin-related aspen genes Potra000605g04596 and Potra002054g16021 (Supplemental Table 2). Arabidopsis mutants for these genes (*HBI1*, *AIF3*, and *AIF4*) have been linked with alteration of petiole length (Supplemental Table 6).

*LMI2* was a highly ranked GR prediction. Importantly, *LMI2* (a MYB TF) is not a paralog of *LATE MERISTEM IDENTITY 1* (*LMI1,* a homeobox TF), also predicted here. Although *LMI1* and *LMI2* belong to different TF families, they both function downstream of LEAFY to regulate meristem transition (Pastore et al., 2011). *LMI1* was reported to regulate leaf growth in Arabidopsis and other species (Vlad et al., 2014; Andres et al., 2017; Li et al., 2021). Arabidopsis *LMI1* loss-of-function mutant showed decreased leaf serration and promoted tissue growth in stipules (Vuolo et al., 2018). The observed phenotype of mutated *LMI2* was related to an increase of the number of cauline leaves and secondary inflorescences (Pastore et al., 2011). Here, *LMI2* transgenic lines were subjected to phenotypic analysis, which demonstrated that a *LMI2* loss-of-function mutant showed increased leaf number and rosette area. We do not exclude that other organs and/or traits might also be affected by the loss of functionality of this gene. The neighborhood conservation of both *LMI1* and *LMI2* suggests that it would be worthwhile to further explore their roles in leaf shape control across monocots and dicots.

Other known examples of functionally conserved predictions across monocots and dicots were GRFs (e.g. the highly ranked *GRF2*), which have a recognized role in leaf size regulation, and AN3/GIF1, a transcriptional co-activator protein (Nelissen et al., 2016). This was also testified by their network conservation in stringent density subnetworks (DS2). A second gene, *GL1*, had its network neighborhood conserved with GRMZM2G022686 from maize. This maize gene encodes for the MYB-related protein *Myb4*. This protein plays important roles in plant improved tolerance to cold and freezing in Arabidopsis and barley (Soltész et al., 2012), but no connections with growth have been observed for this gene. Arabidopsis *SUC2* showed conservation with GRMZM2G307561, a sucrose/H^+^ symporter which remobilize sucrose out of the vacuole to the growing tissues. Mutants for this gene showed reduced growth and the accumulation of large quantities of sugar and starch in vegetative tissues in Arabidopsis (Srivastava et al., 2008), while in maize mutants, slower growth, smaller tassels and ears, and fewer kernels were observed (Leach et al., 2017). This gene is thus also important for growth, development, and yield across monocots and dicots.

The application of a cross-species approach is an important feature of our methodology. To perform GR predictions, translated-GRs from aspen and maize were also used as guide genes, together with triplets to focus on the conserved parts of the inferred leaf networks. As a result, among the cross-species conserved predictions with experimental evidence in more than one species described above, *AUX1*, *IAA3*/*SHY2*, *AXR5*, *AIF3*, *AIF4*, *HBI1*, *AN3/GIF1*, *GL1,* and *SUC2* couldn’t have been predicted using solely primary-GRs from Arabidopsis. This observation indicates that the integration of information of different plant species enhances the detection of GRs.

A total of 11 primary-GRs from Arabidopsis showed no network neighborhood conservation. Lack of conservation might be the result of (1) missing orthologs in a target species or (2) different network gene neighbors across species, which in turn might be caused by different transcriptional control. One clear example of no conservation due to a lack of orthologs is PEAPOD 2 (*PPD2)*, which is a TIFY transcriptional regulator part of the PEAPOD (PPD) pathway. This pathway plays an important role in cell proliferation and, with its PPD/KIX/SAP module, is involved in leaf, flower, fruit, and seed development. This pathway is present in most vascular plant lineages, but was lost in monocot grasses (Schneider et al., 2021). The reason for this absence might be found back in intrinsic differences between eudicots and grasses, being mainly lack of meristemoids and functional redundancy for the regulation of cell proliferation. Surprisingly, several non-grass monocot species such as *Musa acuminata* (banana) and *Elaeis guineensis* (oil palm), basal angiosperm *Amborella trichopoda* and lycophytes, carry PPD/KIX/SAP orthologs, although information about their functionality is missing (Schneider et al., 2021). Another gene with orthologs but lacking network neighborhood conservation was *AHK3*, a cytokinin receptor that controls cytokinin-mediated leaf longevity. This might be explained by knock-out experiments on *AHK* receptors showing contrasting effects on flowering time or floral development across Arabidopsis and rice (Burr et al., 2020). Another non-conserved GR was *ZHD5* that regulates floral architecture and leaf development and is regulated by *MIF1 (MINI ZINC-FINGER 1)* (Hong et al., 2011), which also lacked network conservation. *ZHD5* regulation might thus be different across species. Similarly, FBX*92 (F-BOX PROTEIN92)* was not conserved, which might be explained by the opposite effects on leaf size shown by *ZmFBX92* and *AtFBX92* gain of function in Arabidopsis due to the presence of an F-box-associated domain in *AtFBX92*, lacking in *ZmFBX92*. *FBX92* orthologs might thus undergo different transcriptional regulation (Baute et al., 2017). *EPF1 (EPIDERMAL PATTERNING FACTOR 1)* was also a non-conserved GR. This gene affects stomatal density and water use efficiency. Recent work suggested that, in monocots and dicots, *EPF1* orthologs probably have different temporal dynamics of gene expression in the stomatal lineage (Buckley et al., 2020), which might result in different network gene neighbors.

Based on the validation results of our GR prediction pipeline, a correlation between network size and recovery of genes affecting leaf size was observed. In particular, with increasing network size, the recovery rate decreased, indicating that DS5 is not a recommended network density to use to find new growth regulators. The network neighborhood conservation of genes in the most stringent networks involved different basal biological processes, suggesting their functional similarity across monocots and dicots. Not surprisingly, genes involved in cell cycle regulation and plant hormonal response were found, as both processes have a key role in leaf development. Several cell cycle regulators were predicted as GRs, like the cyclin gene *CYCD3;3*, the *CDK* inhibitor *KRP3 (KIP-RELATED PROTEIN)*, and a DOF transcription factor gene *OBP1* (*OBF BINDING PROTEIN 1*) that controls cell cycle progression (Dewitte et al., 2007; Skirycz et al., 2008; Jun et al., 2013). The auxin-responsive transcription factor gene MONOPTEROS (*MP*) is crucial for leaf vascular development (Hardtke and Berleth, 1998), while the Aux/IAA gene that represses auxin signaling, *AXR2*, whose gain-of-function leads to strong inhibition of leaf growth (Mai et al., 2011), was also predicted. Besides auxin, brassinosteroid (BR) and gibberellin (GA) coordinately play key roles in regulating plant cell elongation. The other two predicted transcription factor genes, *HB25* (*HOMEOBOX PROTEIN 25*) and *MYR1*, which modulate bioactive GA biosynthesis, were also shown to have an effect on the petiole growth (Bueso et al., 2014). It is noteworthy that nearly half of all the 34 genes with leaf phenotype were transcription regulators, which highlights the importance of TF-mediated gene expression regulation during leaf development. In addition to hormone-related genes and TFs, genes related to photosynthesis are also important for leaf development. A carotenoid biosynthesis gene *LCY* and a chloroplast redox-regulating gene *THIOREDOXIN X* were predicted as GR and have been shown to affect leaf size (Li et al., 2009; Pulido et al., 2010). Moreover, the cytoplasmic carbonic anhydrase genes *CA2* and *BCA4* were identified, consistent with the view that carbon utilization in leaves is closely linked to leaf area (DiMario et al., 2016). Cell wall modification is considered to be another important determinant of leaf development. The predicted candidate genes *LACCASE11 (LAC11)* and *CUTICLE DESTRUCTING FACTOR 1* (*CDEF1)*, encoding for a laccase that associates with the lignin deposition in cell wall and a cutinase essential for the degradation of cell wall components, respectively, are also involved in regulating leaf growth and morphology (Takahashi et al., 2010; Qin et al., 2013). Among Arabidopsis genes with a reported phenotype in the RARGE II loss-of-function dataset, *ACO2 (ACC OXIDASE 2)* led to increased leaf size, and AT3G43270, a member of Plant invertase/pectin methylesterase inhibitor superfamily, to smaller leaves. GRs translated from aspen led, through our integrative network approach, to the prediction of *NITRATE TRANSPORTER 1.3* (*NPF6.4*/*NRT1.3)* as a new potential GR. In Arabidopsis shoot, the expression of *AtNPF6.4*/*NRT1.3* was induced by nitrate (Okamoto et al., 2003) while, in *Medicago truncatula*, MtNRT1.3 shares 70% identity with *AtNPF6.4*/*NRT1.3* and was reported to be a dual-affinity nitrate transporter (Morre-Le Paven et al., 2011). It was also hypothesized that *NPF6.4*/*NRT1.3* may play a role in supplying nitrate to photosynthesizing cells (Tong et al., 2016). In our experiments, we showed that this gene, when mutated, is altering leaf growth. This cross-species conserved gene would thus contribute to nitrogen assimilation, that, closely interacting with carbon metabolism, sustains plant growth and development (Nunes-Nesi et al., 2010). Due to the relevance and the strong interconnection of the processes where *NPF6.4/NRT1.3* and many of the candidate GRs here predicted, are involved in, future experimental work will have to reveal the role of these candidate GRs in other organs.

In conclusion, the approach developed in this study fully exploits the potential of integrative biology to translate and expand yield-related functional annotations in different plant species, as such accelerating crop breeding.

## Methods

### Integration of developmental expression datasets and network construction

Transcriptomic datasets were obtained from a list of studies in Arabidopsis, maize and aspen covering samples from the main leaf developmental phases (Supplemental Table 1, Supplemental Methods, Dataset 1). Details about these datasets processing of these samples were reported in Supplemental Table 1. Maize data was mainly composed by a developmental compendium newly generated in this work (Supplemental Methods). The network inference was carried out with Seidr (Schiffthaler et al., 2018), which infers gene networks by using multiple inference algorithms and then aggregating them into a meta-network. This approach has been shown to strongly improve the accuracy of the results (Marbach et al., 2012). Each network was subset into five density subnetworks (DSs) using five different network density values. This procedure consisted in selecting the top 0.1, 0.5, 1, 5 and 10% top Seidr links in each species-specific network and generating five DSs (from the most stringent DS1 to the least stringent DS5).

### Orthology and network neighborhood conservation

To compute cross-species gene network neighborhood conservation, orthology information between genes from Arabidopsis, maize and aspen was computed using the PLAZA comparative genomics platform (Van Bel et al., 2018). A custom version of this platform was built covering in total 15 eukaryotic species including *Arabidopsis thaliana* (TAIR10), *Eucalyptus grandis* (v2.0), *Populus trichocarpa* (v3.01), *Populus tremula* (v1.1), *Vitis vinifera* (12X March 2010 release), *Zea mays* (AGPv3.0), *Oryza sativa* ssp. *Japonica* (MSU RGAP 7), *Triticum aestivum* (TGACv1), *Amborella trichopoda* (Amborella v1.0), *Picea abies* (v1.0), *Pinus taeda* (v1.01), *Selaginella moellendorffii* (v1.0), *Physcomitrella patens* (v3.3), *Chlamydomonas reinhardtii* (v5.5) and *Micromonas commode* (v3.0). PLAZA allows identifying orthologs using different methods (evidences), corresponding to orthologous gene families inferred through sequence-based clustering with OrthoFinder (Emms and Kelly, 2015), phylogenetic trees, and multispecies Best-Hits-and-Inparalogs families (Van Bel et al., 2012). The PLAZA orthology relationships were extracted and filtered retaining all orthologs having a requirement of 2/3 orthology evidences and, for those with 1/3 evidence and >25 orthologs, the ones corresponding to the best 25 blast hits (sorted by e-value) were retained. The generated orthology output was used for the following pipeline steps.

The generated DSs and the orthology information were used to compare the three species using a network neighborhood conservation analysis (ComPlEx analysis, as in Netotea et al.). In this analysis, the network neighborhood of a gene (i.e. all genes with a link to it) was considered conserved if it had a statistically significant (q < 0.05) overlap with the network neighborhood of its ortholog in the other species (Netotea et al., 2014). Here, the comparison was performed for all pairs of networks between the datasets of the three species, and the output of this analysis was collated to create “triplets”. The triplets are sets of three orthologous genes–one per network/species–that have a significantly conserved network neighborhood in all three pairs of comparisons. Since the test is not commutative, the neighborhoods had to be significantly conserved in both directions of the test. To estimate the false discovery rate (FDR) of the detection of triplets, a permutation strategy was adopted. For 500 runs of ComPlEx, ortholog relationships were shuffled, keeping the relative number of orthologs per gene and per species, and then comparing the number of triplets computed from randomization with those resulting using the original (unshuffled) orthologs.

### Functional analyses and prediction of growth regulators

Gene Ontology (Ashburner et al., 2000) functional annotations for Arabidopsis, maize and aspen were retrieved from TAIR (download 25/12/2018), Gramene (AGPv3.30, http://bioinfo.cau.edu.cn/agriGO/download.php), and PlantGenIE (ftp://ftp.plantgenie.org/Data/PopGenIE/Populus_tremula/v1.1/annotation/), respectively, and filtered for the genes present in the corresponding species networks. We focused on biological processes (BP) and excluded the general GO BP terms with >= 1500 genes as well as GO terms with <= 10 genes to avoid biases towards very general and specific terms. For each gene, all GO annotations were recursively propagated in order to include parental GO terms. Functional over-representation analyses were performed using the hypergeometric distribution together with Benjamini-Hochberg (BH) correction for multiple testing (Benjamini and Hochberg, 1995). To get a complete view on all relevant processes related to plant growth, information from literature was collected on growth regulators (GRs). Experimentally validated genes in Arabidopsis, maize and aspen (primary-GRs) were retrieved from public databases (Gonzalez et al., 2010; Beltramino et al., 2018). Experimentally validated aspen genes were obtained by access to SweTree Technologies private database that contains data from the large-scale testing of >1,000 genes and their growth-related properties (here only “stem size” was taken into consideration), an effort where more than 1,500 recombinant DNA constructs were used to either introduce a new gene product or alter the level of an existing gene product by over-expression or RNA interference in aspen trees, whose growth characteristics were then monitored in greenhouse and field experiments to provide extensive gene-to-yield data. The Arabidopsis GR primary set was then enlarged with high quality GR orthologs from maize and aspen using the triplets (“translated-GRs”) to obtain a combined GR set. The combined set was finally filtered with genefilter package for Bioconductor (Gentleman et al., 2021) to remove genes with small expression variance (var.func=IQR, var.cutoff=0.8) and focus on genes active during proliferation or expansion phases of leaf development (“expression-supported GRs”, Supplemental Table 2). Other information on functional categories (Vercruysse et al., 2020a) and differentially expressed genes from relevant studies on plant development was also included in the functional enrichment analyses (Anastasiou et al., 2007; Gonzalez et al., 2010; Eloy et al., 2012; Vercruyssen et al., 2014).

The expression-supported GRs were used as guide genes to perform network-guided gene function prediction via a guilt-by-association (GBA) approach. This approach is based on the assumption that genes close to the input GRs in the network are likely to have similar functions. The GBA approach was applied to attribute new functions based on GO enrichment in the modules of each DS yielding five sets of gene predictions. By this procedure, gene neighborhoods significantly enriched for guide GRs were functionally annotated (hypergeometric distribution). This allowed to predict new GRs and estimate, for each of them, a corresponding FDR adjusted p-value (or q-value), which was renamed “GBA-score”. The GBA score is a confidence score that ranks genes high if they are connected with many GRs in the network (in fact high ranked genes have a low GBA score as this is an indicator of a strong enrichment). For an example GR prediction (in one of any of the five DSs), the GBA-score from the five DSs was summarized taking the mean of the GBA-scores and setting the GBA-score to 0.05 for the DSs where the gene was not predicted. This yielded a list of GR predictions that was then further filtered by only retaining those predictions having conserved neighborhood in at least one DS. To perform a validation of the gene function predictions, the RARGE II (Akiyama et al., 2014) database was interrogated to retrieve a list of Arabidopsis genes that, when mutated, showed an increased or decreased length, width and size for rosette leaf, vascular leaf and cauline leaf (leaf trait genes). This gene set was used to analyze the recovery at each DS of leaf growth-related phenotypes. For the top 100 predictions ranked by GBA-score a manual literature search was performed to retrieve all genes with a reported phenotype including information about the biological pathway the gene might be active in, and other public functional annotations.

### Rosette growth phenotyping

The *Arabidopsis thaliana* ecotype Columbia-0 (Col-0) was used as the wild-type in this study. The T-DNA insertion lines for At4g26530 (Salk_080758/*fba5-1*), At3g21670 (Salk_001553/*sper3-3*), At3g61250 (Salk_066767/*lmi2-1*, Salk_020792/*lmi2-2*), At4g25240 (Salk_113731), At1g63470 (Salk_123590/*ahl5*), At4g37980 (Salk_001773/*chr hpl*), At2g38530 (Salk_026257/*ltp2-1*), At4g28950 (Salk_019272), and At1g12240 (Salk_016136) were confirmed using PCR with a T-DNA primer and gene-specific primers (Lu et al., 2012; Zhao et al., 2013; Jacq et al., 2017; Tanaka et al., 2018; Pastore et al., 2011; Tong et al., 2016). All tested seeds were stratified in the darkness at 4 °C for 3 days and then sown on soil in the 7 cm wide square pots with a density of four seeds per pot. After 8 days in the growth room (with controlled temperature at 22 °C and light intensity 110μmol m^-2^ s^-1^ in a 16 h/8 h cycle), the four seedlings were screened, leaving one seedling per pot, which most closely resembled the genotype average. The plants were imaged in a phenotyping platform (MIRGIS) with fixed cameras located directly above the plants, which images plants at the same time every day. These images were then processed to extract the rosette growth parameters of each plant. The mean PRA, compactness and stockiness values were calculated over time for each genotype.

## Accession Numbers

Sequence data from this article have been submitted to ENA (E-MTAB-11108).

## Supplemental data

Supplemental Figure 1. Number of neighbors per gene at each density subnetwork in Arabidopsis.

Supplemental Figure 2. Expression patterns for the expression-supported growth regulators in Arabidopsis.

Supplemental Figure 3. Expression supported growth regulators with neighborhood conservation at each density level.

Supplemental Figure 4. Functional enrichment of cross-species conserved transcription factors (TF) grouped by TF family.

Supplemental Figure 5. Identification of T-DNA insertion lines.

Supplemental Figure 6. The rosette leaf numbers of the wild-type Col-0 and the mutants of *NRT1.3* and *LMI2*.

Supplemental Table 1. Overview of the expression datasets used for the network computation

Supplemental Table 2. List of expression-supported growth regulators

Supplemental Table 3. Predicted growth regulators

Supplemental Table 4. List of RARGE II leaf trait genes known to affect leaf phenotype if mutated

Supplemental Table 5. Top 100 predicted growth regulators annotated

Supplemental Table 6. In depth literature analysis for the top 100 predicted growth regulators

Supplemental Table 7. List of genes tested for leaf phenotype in this study

Supplemental Dataset 1. Expression datasets for Arabidopsis, maize, and aspen

Supplemental Dataset 2. Triplets generated with ComPlEx

Supplemental Methods. Detailed methods for expression dataset retrieval, generation, and processing.

## Acknowledgements

We thank Taku Takahashi (Okayama University, Japan) for kindly sending us the *sper3-1* and *sper3-3* seeds, and Julie Pevernagie for her technical support.

