## Supplemental information for "Identification of new growth regulators using cross-species network analysis in plants"

### Supplemental Methods

### Maize developmental expression dataset

#### Maize growth conditions

Maize plants were grown in growth chambers with controlled relative humidity (55%), temperature (24 °C day/18 °C night), and light intensity (170–200 µmol $m^{-2}$ $s^{-1}$ photosynthetic active radiation at plant level) provided by a combination of high-pressure sodium vapor (RNP-T/LR/400W/S/230/E40; Radium) and metal halide lamps with quartz burners (HRI-BT/400W/D230/E40; Radium) in a 16-h/8-h (day/night) cycle.

#### Developmental maize compendium (15 samples)

Three sections (from the base to 3.5 cm, from 3.5 to 7.0 cm and from 7.0 to 10.5 cm) of a developing leaf 4 were harvested two days after leaf emergence, from maize B104 inbred plants. To aim for enough tissue per section and per replicate, 28 plants per replicate were pooled. In total, five biological replicates for the three sections (15 samples in total) were used for RNAseq. After harvesting, samples were directly frozen in liquid nitrogen. Total RNA was extracted using the guanidinium thiocyanate-phenol-chloroform extraction method using TRI-reagent (Thermo Fisher Scientific) followed by DNA digestion using the RQ1 RNase-free DNase kit (Promega). Total RNA was sent to GATC Biotech for RNA sequencing. Library preparation was done using the NEBNext Kit (Illumina). In brief, purified poly(A)-containing mRNA molecules were fragmented, randomly primed strand-specific cDNA was generated and adapters were ligated. After quality control using an Advanced Analytical Technologies Fragment Analyzer, clusters were generated through amplification using cBOT (Cluster Kit v4, Illumina), followed by sequencing on an Illumina Hi Seq2500 with the TruSeq SBS Kit v3 (Illumina). Sequencing was performed in paired-end mode with a read length of 125 nt.

#### qRT-PCR for zone delineation in the developmental maize compendium (methods)

The first ten cm of a growing fourth leaf, two days after leaf emergence, from maize B104 inbred lines was harvested and segmented into smaller pieces of 5mm (basal two cm) and 10mm (distal eight cm). For each piece, we had three biological replicates, each pool consisting of tissue of three plants. After harvesting, samples were directly frozen in liquid nitrogen. Total RNA was extracted using the guanidinium thiocyanate-phenol-chloroform extraction method using TRI-reagent

(Thermo Fisher Scientific) followed by DNA digestion using the RQ1 RNase-free DNase kit (Promega). cDNA was prepared from 1 µg of total RNA with the iScript cDNA Synthesis Kit (Biorad). The qRT-PCR was done on a Lightcycler 480 (Roche) with SYBR green for detection in a 5-µl volume (2,5 µl of mastermix, 0,25 µl of 5 µM of each forward and reverse primer and 2 µl of cDNA). Every reaction was performed in triplicate on a 384-multiwell plate to allow determination of mean and SEM of cycle threshold (CT) values. Data were analyzed in Microsoft Excel with the 2-ΔΔCT method (Schmittgen and Livak, 2008) and values were standardized against those of 18S rRNA (primers P1 and P2). The mean expression levels were calculated from three biological repeats, using the P3 and P4 primers for phosphoribulokinase, P5 and P6 for NADP malate dehydrogenase, P7 and P8 for NADP-malic enzyme (NADP-ME), P9 and P10 for Photosystem Q(B) protein (psbA), P11 and P12 for cytochrome B6 (petB), P13 and P14 for NADPH-quinone oxidoreductase subunit 1 (ndhA), P15 and P16 for Photosystem I iron-sulfur center (psaC) and P17 and P18 for phosphoenolpyruvate carboxylase (PEPC) (**Supplemental methods figure 1**).

**qRT-PCR for zone delineation in the developmental maize compendium (assay results)**

Throughout the developmental gradient represented in the maize leaf growth zone, genes related to photosynthesis are differentially expressed, some even starting in the division zone and expansion zone (Nelissen et al., 2018). Therefore, the maize RNAseq compendium along the developmental gradient of a growing maize leaf three zones were delineated based on a qRT-PCR analysis of several known genes involved in photosynthesis (Wang et al., 2014; Chotewutmontri and Barkan, 2016; Schlüter and Weber, 2019; Heldt and Piechulla, 2021). The qRT-PCR results showed that those genes had specific transcriptional profiles in the lower half of maize leaves that can be divided in three classes. The fragment from the base to 3.5 cm, contains the leaf growth zone in which only the tested transcripts involved in the light dependent reactions of photosystem I and II (Photosystem Q(B) protein (psbA), cytochrome B6 (petB), NADPH-quinone oxidoreductase subunit 1 (ndhA) and Photosystem I iron-sulfur center (psaC)) were expressed (**Supplemental methods figure 1**). Their expression gradually increased along the leaf developmental gradient. The expression level of the other tested genes involved in the C4 carbon assimilation cycle was minimal at the base of the leaf and their transcription levels started to increase from 3.5 to 7.0 cm (NADP malate dehydrogenase and phosphoenolpyruvate carboxylase (PEPC)) or even only started to show an increase in expression in the mature part of the leaf from

7.0 to 10.5 cm (NADP-malic enzyme (NADP-ME) and phosphoribulokinase) (**Supplemental methods figure 1**). From 7.0 to 10.5 cm the expression of all genes had reached their maximal value.

In conclusion, based on a qRT-PCR analysis of genes involved in photosynthesis, three zones along the developmental gradient of the maize leaf were harvested. While the first section consisted of proliferative and expanding leaf tissue (base to 3.5 cm), the second section (3.5 to 7.0 cm) contained expanding and mature cells and the last part (7.0 to 10.5 cm) was fully mature.

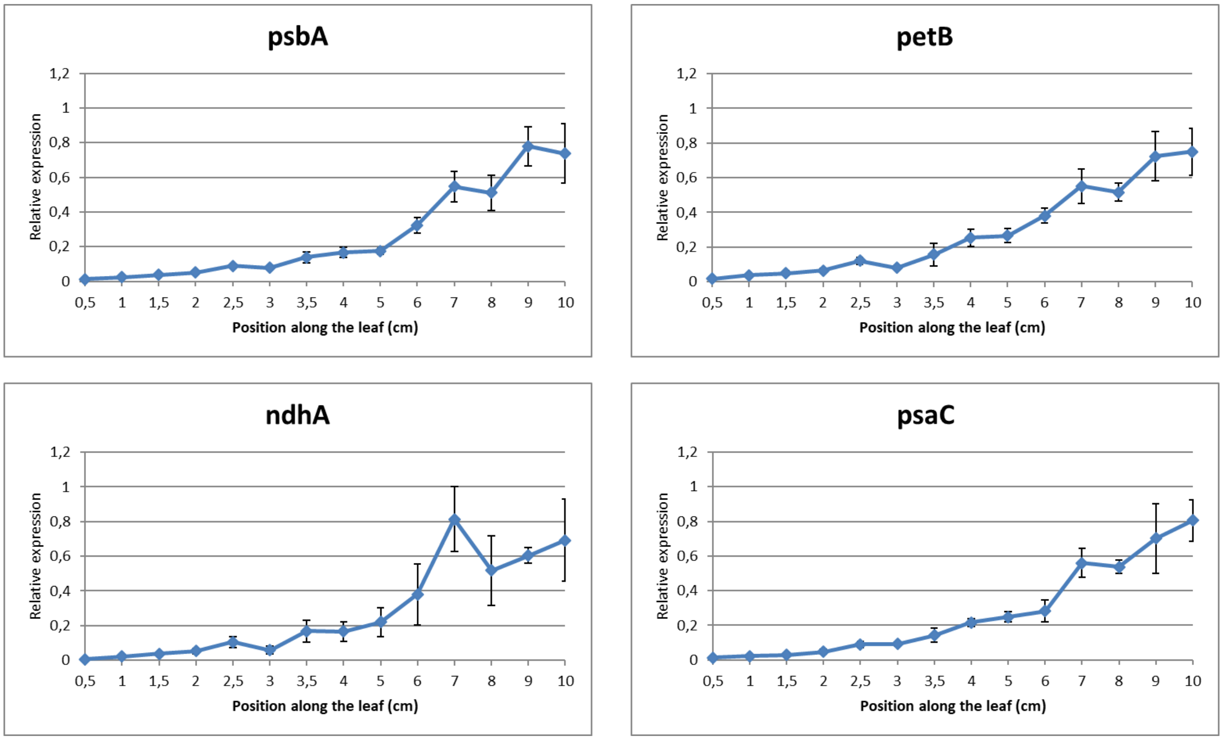

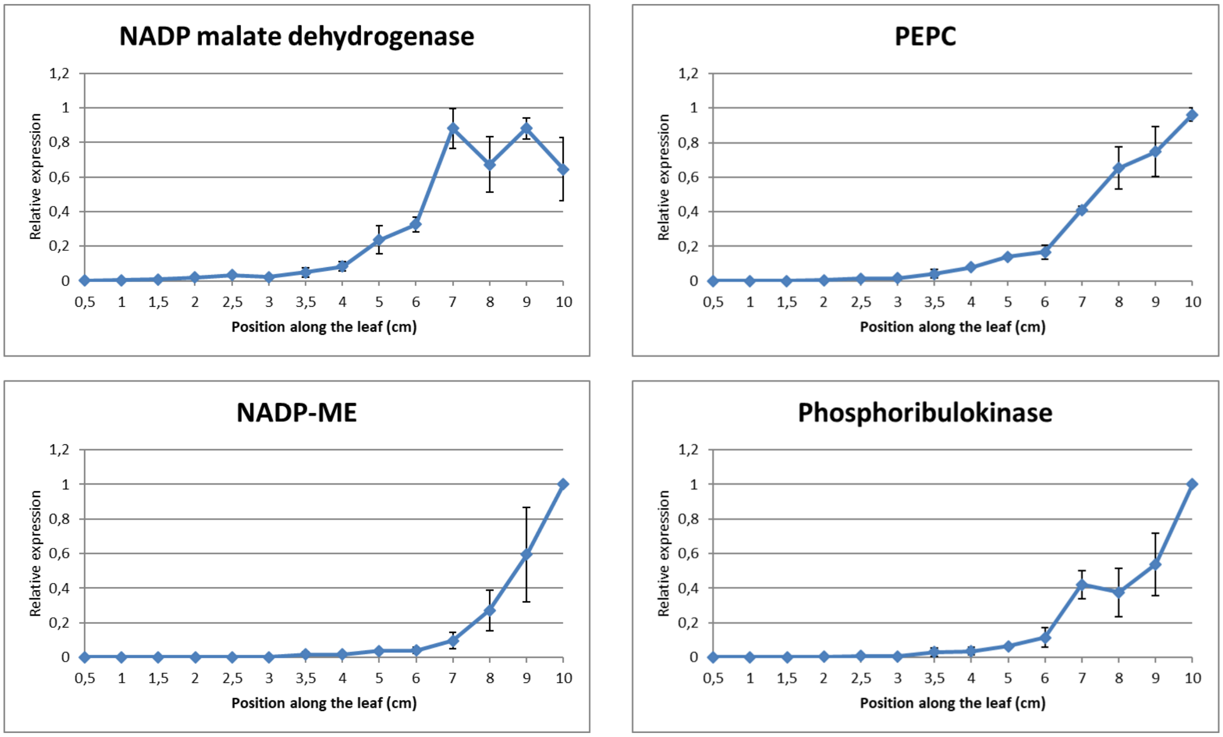


**Supplemental methods figure 1.** Transcripts encoding critical C4 photosynthesis enzymes are differentially expressed in a gradient fashion in the lower half of B104 maize leaves.

#### Proliferative maize samples (3 samples)

The three proliferative maize dataset samples were taken from the inbred line B104. The first basal half cm (dividing cells) of leaf four two days after leaf appearance was sampled. Three biological replicates were taken, each pool consisting of proliferative tissue of three plants. After harvesting, samples were directly frozen in liquid nitrogen. Total RNA was extracted using the guanidinium thiocyanate-phenol-chloroform extraction method using TRI-reagent (Sigma-Aldrich). RNA concentration and purity were determined spectrophotometrically using the Nanodrop ND-1000 (Nanodrop Technologies) and RNA integrity was assessed using a Bioanalyser 2100 (Agilent). Per sample, 500 ng of total RNA was used as input. Using the Illumina TruSeq® Stranded mRNA Sample Prep Kit (protocol 15031047 Rev E October 2013) poly-A containing mRNA molecules were purified from the total RNA input using poly-T oligo-attached magnetic beads. In a reverse transcription reaction using random primers, RNA was converted into first strand cDNA and subsequently converted into double-stranded cDNA in a second strand cDNA synthesis reaction. The cDNA fragments were extended with a single 'A' base to the 3' ends of the blunt-ended cDNA fragments after which multiple indexing adapters were ligated introducing different barcodes for each sample. Finally, enrichment PCR was carried out to enrich those DNA fragments that have adapter molecules on both ends and to amplify the amount of DNA in the library. For the sequence run, libraries were equimolarly pooled and sequenced using a high 300 cycles (PE- 2 x 150 bp) NextSeq kit. Sequencing was performed on an Illumina NextSeq 500 Paired-End mode.

**Other maize samples (6 samples)**

Other six maize samples corresponding to proliferation stage of developing leaf 4 were obtained from Sun et al. (2017) (see the original article for more details).

**Maize data processing**

The 24 total RNA-seq sample reads were processed with Prose (Vaneechoutte and Vandepoele, 2019), which implements kallisto (Bray et al., 2016) for mapping against the maize genome version B73 RefGen_v3.

**Aspen developmental expression dataset (see the original article for more details)**

Aspen data was obtained by the developmental series of terminal leaves published by (Mähler et al., 2020) (LeafDev dataset, 33 samples). This dataset was composed by: the first fully unfurled leaf, defined as a reference point and labeled leaf T0; three leaves above the reference leaf (labeled as T-1, T-2, and T-3) and the apical region, containing the shoot apical meristem; the very youngest leaf primordia (labeled T-4); and two leaves below the reference leaf (labeled T1 and T2).

**Arabidopsis developmental expression dataset (see the original articles for more details)**

Transcriptomic data for Arabidopsis were obtained from several studies: AGRONOMICS1 Tiling Array (Andriankaja et al., 2012) including leaves from seedlings harvested at the stages of proliferation (8 and 9 days after sowing (DAS)), transition (10, 11, and 12 DAS), and expansion (13 and 14 DAS) for a total of 24 samples; ATH1-array (Skirycz et al., 2010) including leaves harvested from plants at proliferation (9 DAS) and expansion (15 DAS) stages for a total of 6 samples. ATH1-array (Skirycz et al., 2011) including leaves harvested at proliferation stage (9DAS). RNA-seq data (Dubois et al., 2017) including leaves harvested at expansion stage (11 DAS) for a total of 11 samples.

**Arabidopsis data processing**

The integration of array and RNA-seq data followed two main steps. The first was performed to obtain two datasets with the same distribution and was performed via the quantile normalization selecting processed RNA-seq data sample-set (11 samples) as target distribution and microarray sample-set (42 samples) as reference distribution (Thompson et al., 2016). In the second step, all samples were inspected using principal component analysis (PCA). Batch effect correction was applied using ComBat implemented in the R package SVA to remove non-biological sources of variation in the dataset (Leek et al., 2010).

### Supplemental Figures: Identification of new growth regulators using cross-species network analysis in plants

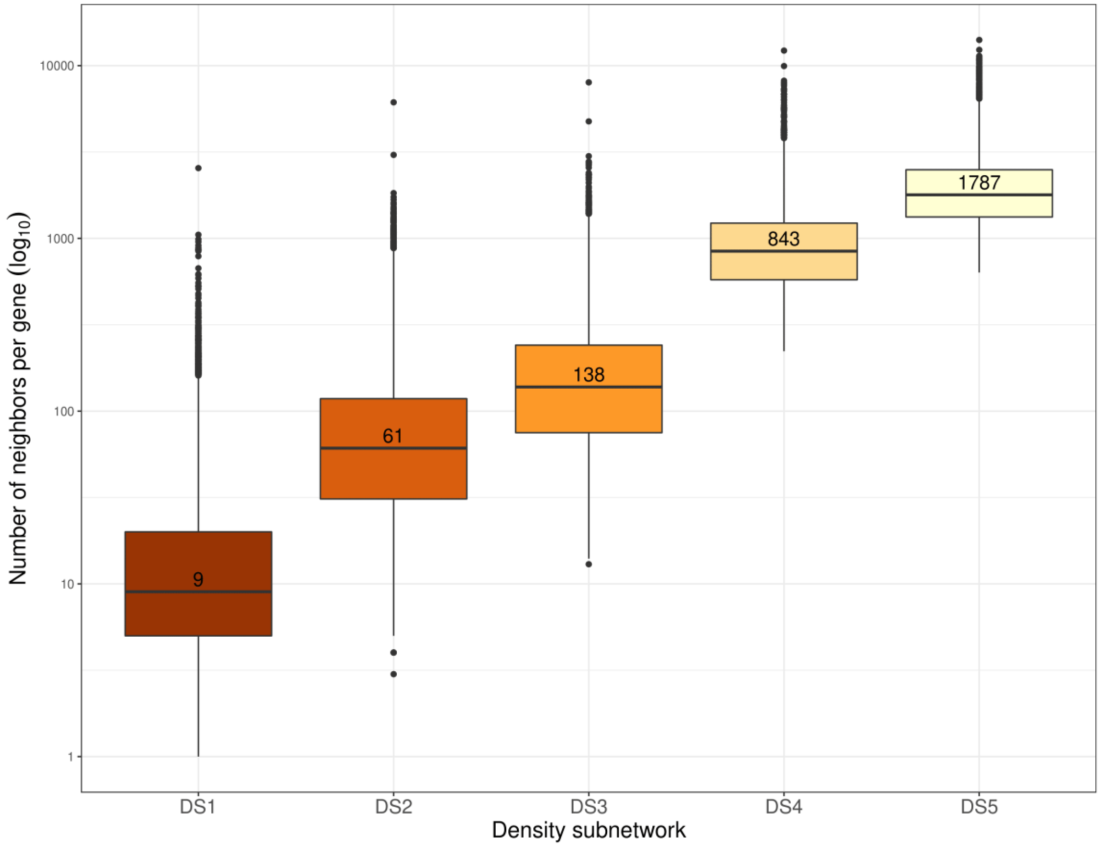


**Supplemental Figure 1. Number of neighbours per gene at each density subnetwork in Arabidopsis.**

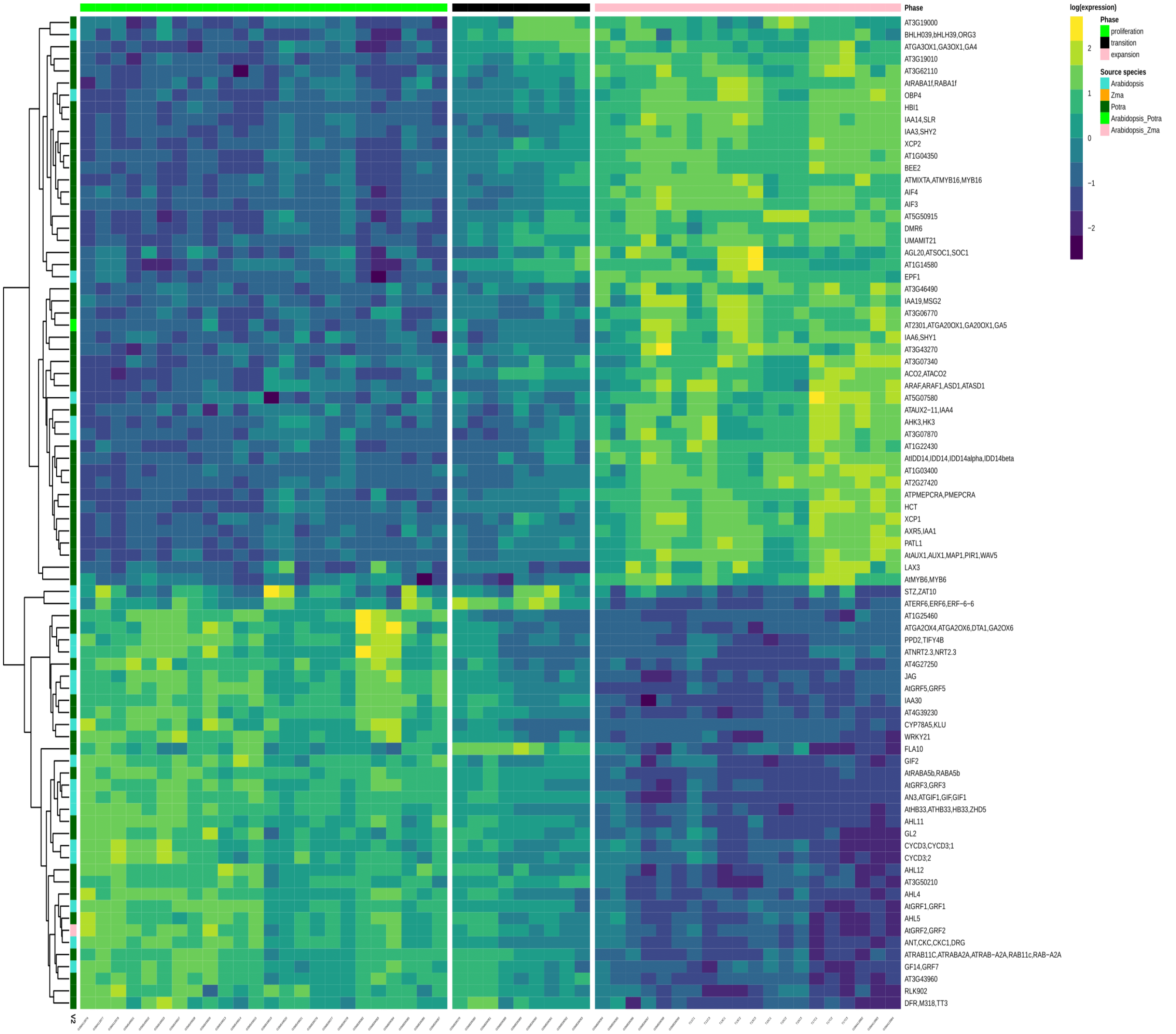


**Supplemental Figure 2**. **Expression patterns for the expression supported growth regulators in Arabidopsis**. Growth regulator sources are also presented (Arabidopsis, aspen, maize, or shared across two species). Values are row-scaled.

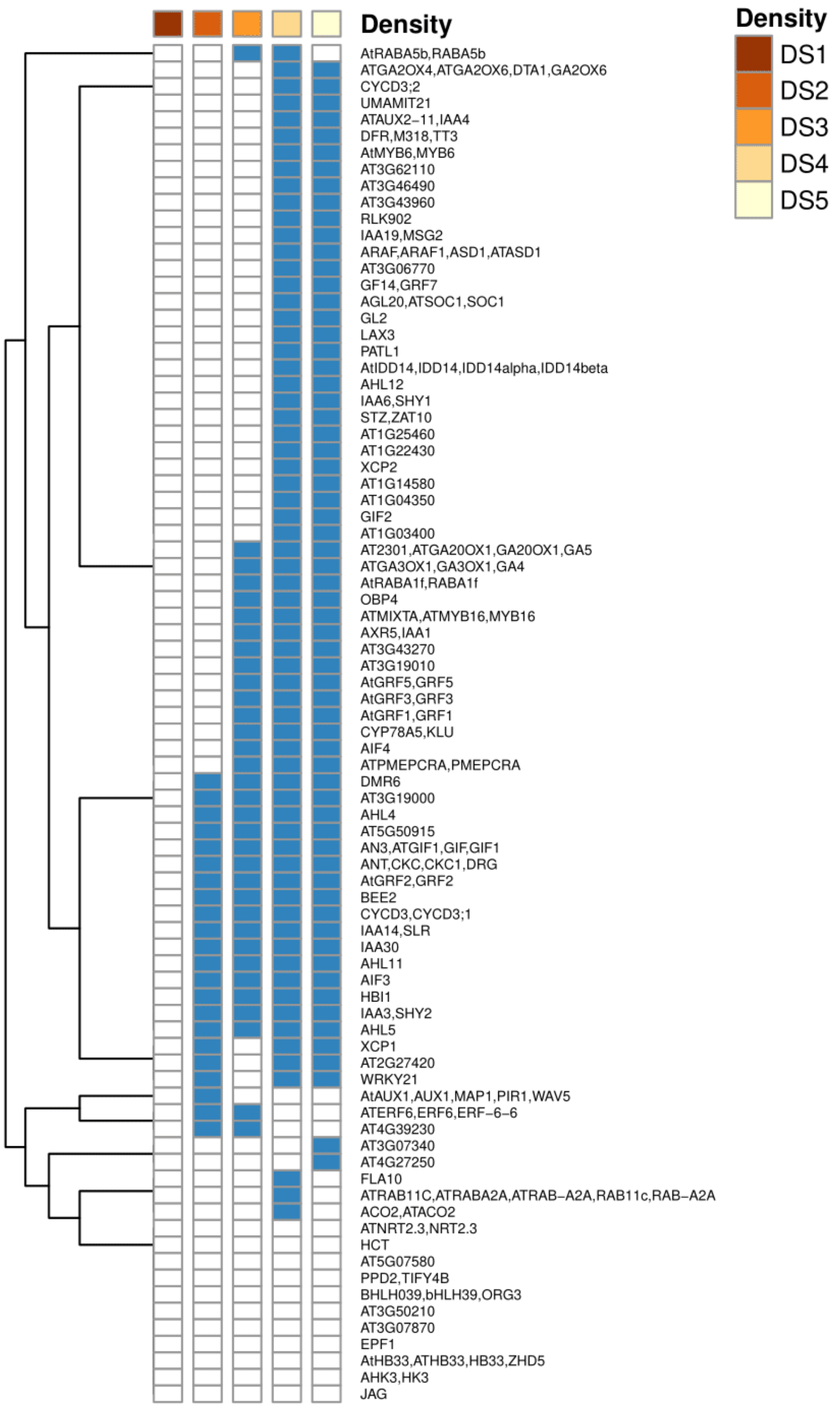


**Supplemental Figure 3. Expression supported growth regulators with neighborhood conservation at each network density level.**

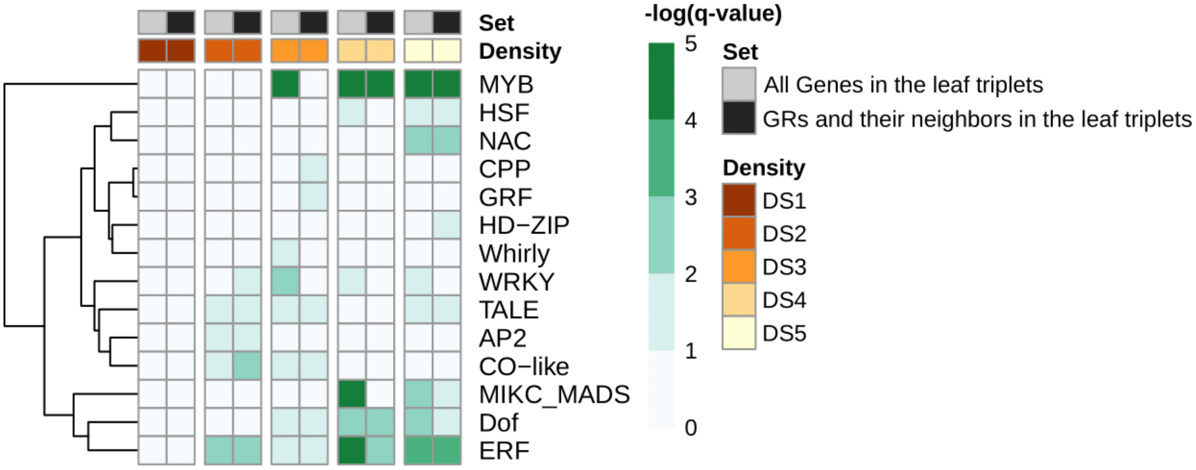


**Supplemental Figure 4**. **Functional enrichment of cross-species conserved transcription factors (TF) grouped by TF family**. Values are expressed as –log(q-value) resulting from the enrichment analysis.

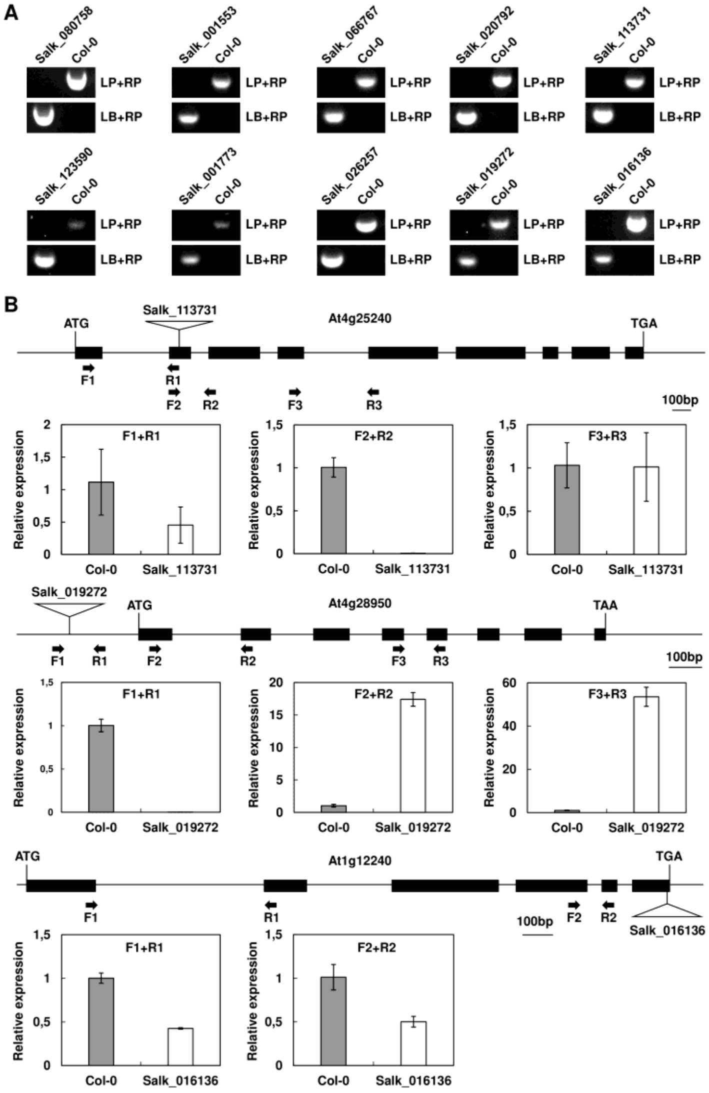


**Supplemental Figure 5. Identification of T-DNA insertion lines.** (A) Molecular analysis of T-DNA insertion lines by PCR using a T-DNA primer and gene-specific primers. (B) qRT-PCR analysis showed the disrupted expression of At4g25240 in Salk_113731, the increased expression of At4g28950 in Salk_019272, and the decreased expression of At1g12240 in Salk_016136, respectively.

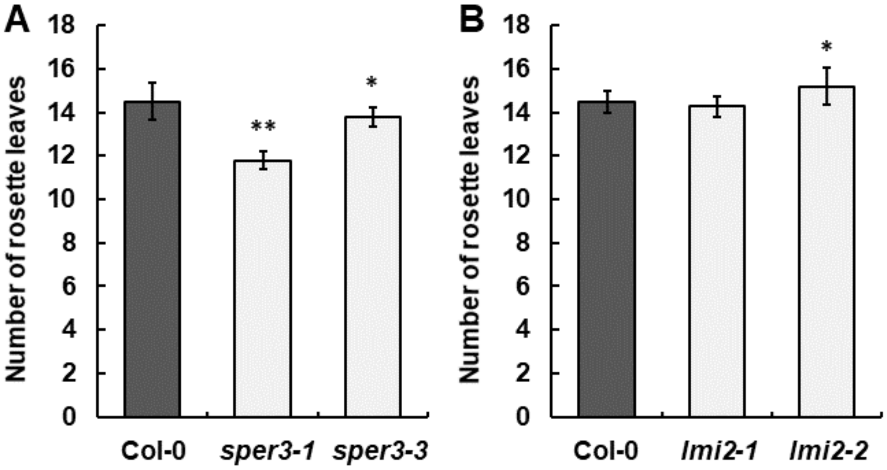


**Supplemental Figure 6. The rosette leaf numbers of the wild-type Col-0 and the mutants of *NRT1.3* and *LMI2*.** The rosette leaf number of 26-day-old wild-type Col-0 and the mutants of *NRT1.3* (A) and *LMI2* (B). Asterisks denote significant differences compared to the wild-type Col-0, as determined by Student's *t* test (*, $P$<0.05; **, $P$<0.01).
